# Biochemical identification of a nuclear coactivator protein required for AtrR-dependent gene regulation in *Aspergillus fumigatus*

**DOI:** 10.1101/2022.09.26.509624

**Authors:** Sanjoy Paul, Shivani Ror, W. Hayes McDonald, W. Scott Moye-Rowley

## Abstract

Azole drugs represent the primary means of treating infections associated with the filamentous fungal pathogen *Aspergillus fumigatus*. A central player in azole resistance is the Zn_2_Cys_6_ zinc cluster-containing transcription factor AtrR. This factor stimulates expression of the both the *cyp51A* gene encoding the azole drug target enzyme as well as an ATP-binding cassette transporter-encoding gene called *abcG1* (aka *cdr1B*). We have used a fusion protein between AtrR and the tandem affinity purification (TAP) moiety to purify proteins that associate with AtrR from *A. fumigatus*. Protein fractions associated with AtrR-TAP were subjected to MudPIT mass spectrometry and one of the proteins identified was encoded by the *AFUA_6g08010* gene. We have designated this protein NcaA (<u>N</u>uclear <u>C</u>o-Activator of <u>A</u>trR). Loss of *ncaA* caused a reduction in voriconazole resistance and drug-induced *abcG1* expression, although did not impact induction of *cyp51A* transcription. We confirmed association of AtrR and NcaA by co-immunoprecipitation from otherwise wild-type cells. Expression of fusion proteins between AtrR and NcaA with green fluorescent protein allowed determination that these two proteins were localized in the *A. fumigatus* nucleus. Together, these data support the view that NcaA is required for nuclear gene transcription controlled by AtrR.

**Importance:** *Aspergillus fumigatus* is the major filamentous fungal pathogen in humans and is susceptible to the azole antifungal class of drugs. However, loss of azole susceptibility has been detected with increasing frequency in the clinic and infections associated with these azole resistant isolates linked to treatment failure and worse outcomes. Many of these azole resistant mutant strains contain mutant alleles of the *cyp51A* gene encoding the azole drug target. A transcription factor essential for *cyp51A* gene transcription has been identified and designated AtrR. AtrR is required for azole inducible *cyp51A* transcription but we know little of the regulation of this transcription factor. Using a biochemical approach, we identify a new protein called NcaA that is involved in regulation of AtrR at certain target gene promoters. Understanding the mechanisms controlling AtrR function is an important goal in preventing or reversing azole resistance in this pathogen.

## Introduction

*Aspergillus fumigatus* is the primary filamentous fungal pathogen of humans and treatment of infections associated with this fungus is complicated due to problems with both diagnosis and limited antifungal therapies (discussed in (1, 2)). A major complication arising with a troubling frequency is the appearance of azole-resistant *A. fumigatus* associated with aspergillosis (reviewed in (3)). Infections caused by *A. fumigatus* strains that are azole resistant have a significantly increased rate of mortality (4), making the understanding of mechanisms underlying azole resistance a high priority.

The best characterized mechanism of azole resistance in *A. fumigatus* involves linked changes in the gene producing the enzymatic target of azole drugs in this organism. This locus is called *cyp51A* and directs the production of the lanosterol α-14 demethylase enzyme, an essential step in the biosynthesis of ergosterol (5). Substitution mutations in the coding sequence of *cyp51A*, along with duplications in the promoter region of this gene, trigger a large decrease in azole susceptibility (6-8). These promoter duplications elicit increased levels of *cyp51A* gene transcription and are required for the relevant clinical phenotypes to be observed (9, 10). This critical link between transcription and drug resistance places a premium on analysis of the mechanisms underlying regulation of *cyp51A*

Findings from several groups have implicated the *A. fumigatus* transcription factors SrbA and AtrR as essential contributors to expression of *cyp51A* (11-13). SrbA was discovered as a key regulator of gene expression in the ergosterol pathway based on its similarity to mammalian SREBP which serves a similar function in control of cholesterol biosynthesis (14). AtrR was initially detected as a positive regulator of expression of ATP-binding cassette (ABC) transporter-encoding gene expression and later shown to be important in transcription of both *cyp51A* as well as the ABC transporter-encoding locus *abcG1* (aka *cdr1B*/*abcC*) in *A. fumigatus* (12, 13).

While basic features of AtrR-responsive gene expression have been described, essentially nothing is known of how this factor is regulated, unlike the case for SrbA (15). To begin to investigate the modulators of AtrR transcription factor function, we prepared a functional fusion protein consisting of full-length AtrR fused at its C-terminus to the tandem affinity purification (TAP) moiety. This AtrR-TAP fusion protein was expressed either from the native *atrR* promoter or from the strong hspA promoter. We prepared highly purified AtrR-TAP under gentle isolation conditions and analyzed the spectrum of co-purifying proteins using Multidimensional Protein Identification Technology (MudPIT) (16). Several proteins were identified that co-purified with AtrR-TAP and here we describe the characterization of a novel nuclear protein we have designated nuclear co-regulation of AtrR (NcaA).

## Materials and Methods

### *A. fumigatus* strains, growth conditions, and transformation

The list of strains that were used in this study are listed in Table 1. *A. fumigatus* strains were routinely grown at 37°C in rich medium (Sabouraud dextrose; 0.5% tryptone, 0.5% peptone, 2% dextrose [pH 5.6 ± 0.2]). Selection of transformants were made in minimal medium (MM; 1% glucose, nitrate salts, trace elements, 2% agar [pH 6.5]); trace elements, vitamins, and nitrate salts are as described in the appendix of reference (17), supplemented with 1% sorbitol and either 20 mg/liter phleomycin (after adjusting the pH to 7) or 150 mg/liter hygromycin Gold (both InvivoGen). For solid medium, 1.5% agar was added. Dox off promoter shut off experiments were performed by adding 25 mg/liter doxycycline (BD Biosciences).

**TABLE 1.**
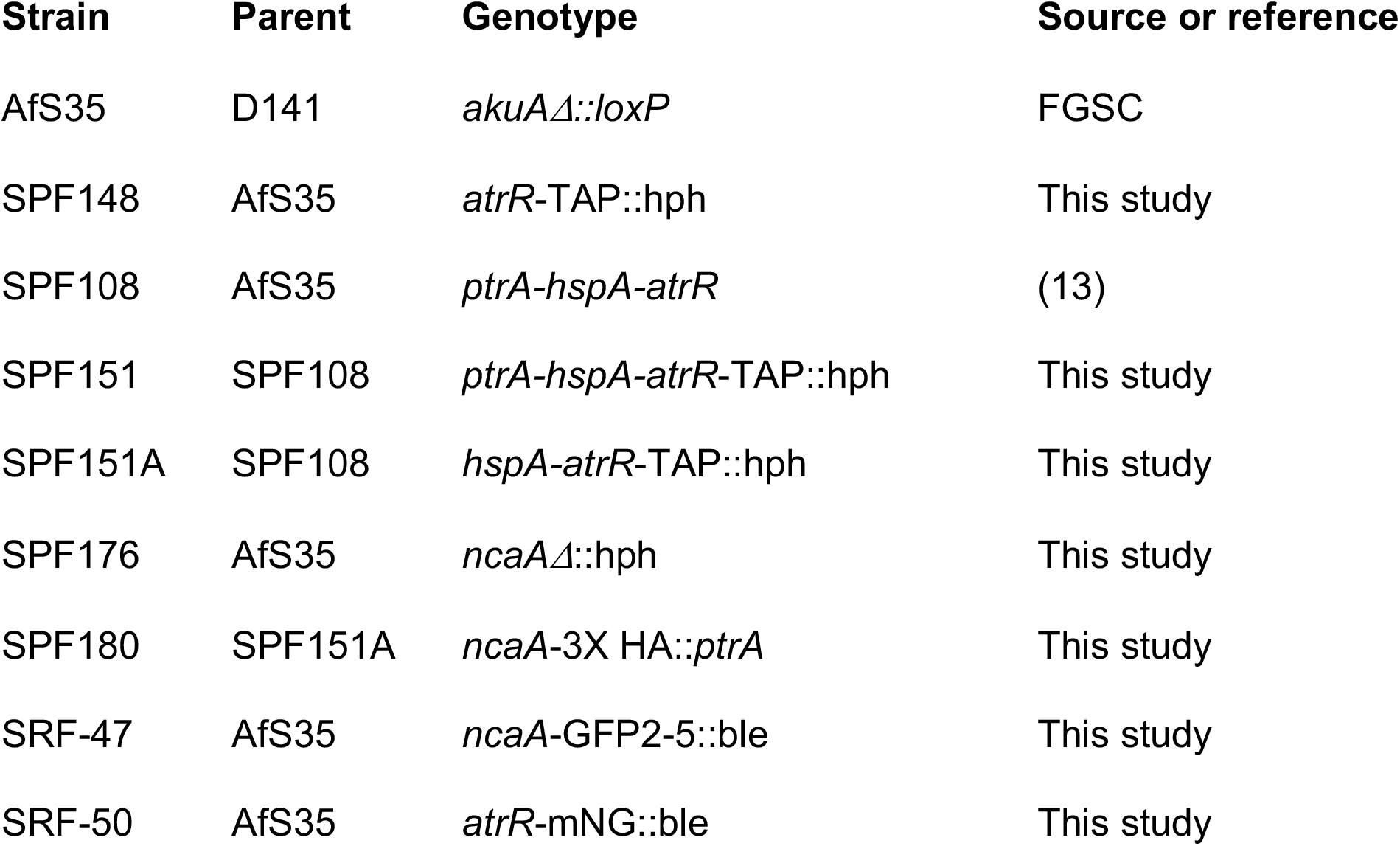
*A. fumigatus* strains used in this study

| Strain | Parent | Genotype | Source or reference |
| --- | --- | --- | --- |
| AfS35 | D141 | *akuAΔ::loxP* | FGSC |
| SPF148 | AfS35 | *atrR-TAP::hph* | This study |
| SPF108 | AfS35 | *ptrA-hspA-atrR* | (13) |
| SPF151 | SPF108 | *ptrA-hspA-atrR-TAP::hph* | This study |
| SPF151A | SPF108 | *hspA-atrR-TAP::hph* | This study |
| SPF176 | AfS35 | *ncaAΔ::hph* | This study |
| SPF180 | SPF151A | *ncaA-3X HA::ptrA* | This study |
| SRF-47 | AfS35 | *ncaA-GFP2-5::ble* | This study |
| SRF-50 | AfS35 | *atrR-mNG::ble* | This study |

Generation of *atrR*-TAP tagged strain was done as follows. Plasmid pSP110 was constructed using Gibson assembly of 4 PCR fragments in pUC19 vector: 1.2 kb corresponding to the 3’ end of *atrR* gene, the G5 linker-TAP tag from plasmid pME4543 (from Bastian Joehnk and Gerhard Braus, see (18) for description of the codon-optimized TAP cassette), the transcription terminator-hph cassette from pSP98 (12), and 1.2 kb region downstream of *atrR* gene. To ensure accurate construction, pSP110 was sequenced to verify the integrity of the 3’ region of *atrR* gene-G5 linker-TAP fusion present in the plasmid. This plasmid was cut with KpnI and HindIII restriction enzymes and transformed into either AfS35 or the SPF108 strain to generate *atrR*-TAP and *hspA-atrR*-TAP strains, respectively. Targeted integration of these strains was verified by PCR diagnosis of novel junctions formed downstream of the TAP tag as well as by Western blotting of the strains using both AtrR and TAP antibodies. Transformation and generation of *ncaA*Δ mutants was done using in vitro-assembled cas9-guide RNA-ribonucleoproteins coupled with 50bp microhomology repair templates (19). For deletion of *ncaA*, CRISPR RNAs (5’-CAGCTGTGACGCACAAGCGCGGG) corresponding to 5’ end of the gene and (5’-TACGCCCCAGAGCTAGGCGGTGG) corresponding to 3’ end of the gene were used to replace *ncaA* with the hygromycin resistance marker cassette amplified from the plasmid pSP62 (13) using ultramer grade oligonucleotides from IDT (primer pairs ncaA-MH-Hph-F: 5’-TGCAATCTCAGGCCCCACTCTTCATCGTTCCAGCTGTGACGCACAAGCGCcctctaaacaagtgtacctgtg and ncaA-MH-Hph-R: 5’-CGGCGGCTATCCGGTTTGGTAAGATCAAACTACGCCCCAGAGCTAGGCGGgatagctctgtacagtgaccg) harboring 50 bp homology to the *ncaA* gene. The *ncaA*-GA5 linker-3X HA tagged strain was generated using CRISPR RNA (5’-TACGCCCCAGAGCTAGGCGGTGG corresponding to 3’ end of the gene) to C-terminally tag *ncaA* with the 3X HA-pyrithiamine resistance marker cassette amplified from the plasmid pSP102 using ultramer grade oligonucleotides from IDT (primer pairs ncaA-MH-ptrA-F: 5’-GCCACCAGTCCCTTCCGGGTTCAAACTTGGCTAGTGCGTCTGGACCACCGCCTAGCTCTGGGGCGggtgctggtgccggt and ncaA-MH-ptrA-R: 5’-TATGTAGGCTGATGGCGGCGGCTATCCGGTTTGGTAAGATCAAACTACGCatctgacagacgggcaattg) harboring 50 bp homology to insert the GA5 linker-3X HA in place of the *ncaA* gene stop codon. The *ncaA-*GFP2-5 locus was constructed by using the pSR35 vector which contains a codon-optimized GFP2-5 gene (20) for expression in *A. fumigatus*. The PCR amplicon obtained with oligos ncaA-GFP2-5 MH F (CAGTCCCTTCCGGGTTCAA ACTTGGCTAGTGCGTCTGGACCACCGCCTAGCTCTGGGGCGGGAGCCGGTGCCA TGCC) and ncaA-GFP2-5 MH R (TCCTGAGGCCTACATATGTAGGCTGATGGCGGCGGCTA TCCGGTCAGTCCTGCTCCTCGGC), which contains homologous regions before and after the stop codon of *ncaA*, the GFP2-5 tag and phleomycin resistance gene. The AtrR protein was C-terminally tagged with *A. fumigatus*-optimized fluorescent protein mNeonGreen by using vector pSR25 (synthesized by GenScript, Piscataway, NJ). Primer pairs AtrR-CoNG MH F (CCCGGTCTTCGACACCA ATGGTCCACCCCACGGTGGATTGGCTGGTGCCGGTGCTGGT) and AtrR-CoNG MH R (GCCCAAATAAGCCTCCCACGCTGGTGTCCGATTCGTTATTTCAGTCCTGCTCCTCGC) were utilized for amplification of a PCR product corresponding to a homologous egion across the *atrR* stop codon, along with the mNeonGreen tag and phleomycin resistance marker gene. All the above C-terminally tagged strains were generated in the AfS35 background and genotypically confirmed by diagnostic PCR of the novel upstream and downstream junction formed upon targeted integration. At least 3 independently targeted transformants were phenotyped for all *ncaA* mutants, of which a representative strain is depicted in the data presented. The list of strains used in this study is given in Table 1.

### TAP purification

AtrR was purified from *atrR*-TAP and *hspA-atrR*-TAP strains based on the protocol derived from (21). Approximately 10^6^ spores of the TAP-tagged strains were inoculated in petri dishes containing 20 ml of Sabouraud dextrose broth at 37°C for 24 h. Mycelia that formed as a biofilm on the top were collected (∼5 g) and ground into fine powder in liquid nitrogen using a mortar and pestle. The ground mycelium was resuspended in 10 ml B250 buffer [250mM NaCl, 100mM Tris-HCl pH 7.5, 0.1% NP-40, 10% glycerol, 1mM EDTA, 1mM DTT, 1mM PMSF, as well as 200 ml of Protease Inhibitor Cocktail for use with fungal and yeast extracts (Sigma, P8215)] in ice cold SS34 tubes, and incubated for 10 min at 4°C, with intermittent vortexing of 30 seconds at setting 6 (Vortex Genie 2 Scientific Industries) every 2 min. The tubes containing resuspended mycelia was centrifuged at 25000 x g for 30 min at 4°C. The supernatant was transferred to ice cold 15 ml tubes containing 1ml of IgG Sepharose 6 Fast Flow (GE Healthcare) and incubated on a rotating platform for 3 hours at 4°C. The crude extract/IgG sepharose suspension was poured into chromatography columns and the extract allowed to flow through with gravity. The IgG Sepharose was then washed twice with 10 ml W250 buffer [250mM NaCl, 40mM Tris-HCl pH 7.5, 0.1% NP-40, 1mM DTT, 1mM PMSF, as well as 100 μl of Protease Inhibitor Cocktail for use with fungal and yeast extracts (Sigma, P8215)], once with 10 ml W150 buffer [150mM NaCl, 40mM Tris-HCl pH 7.5, 0.1% NP-40, 1mM DTT], and finally once with TEV cleavage buffer (TCB) [W150 buffer + 0.5 mM EDTA]. After the TCB wash, the chromatography column was resuspended with 1 ml TCB as well as 20 μl (200 U) of TE protease enzyme (Genscript, Inc) on a rotator and the columns incubated at 4°C for 16 hours. The TEV protease treated suspension was then transferred into the new columns containing 500 μl Calmodulin Sepharose 4B (GE Healthcare) and 6 ml of CBB buffer [150 mM NaCl, 40 mM Tris–HCl pH 8.0,1 mM MgOAc, 2 mM CaCl_2_, 100 mM imidazole, 10mM β-mercaptoethanol] and incubated on a rotating platform at 4°C for 1 h. At the end of incubation, the CBB was allowed to flow through. The column was then washed three times with 1 ml CBB (containing 0.02% NP-40). The proteins were extracted from the Calmodulin column by adding 1 ml EB buffer [W150 + 20mM EGTA + 1 mM MgOAc + 0.02% NP-40 + 10mM β-mercaptoethanol] into the columns, and incubating for 5 min at room temperature. The elution step was repeated again with 1 ml EB. The 2 ml eluate was split into 2x 750ml (for mass spectrometry analysis) and 1x 500ml (for Coomassie/silver staining), precipitated in 25% TCA, and finally the pellet was washed with 1 ml cold acetone. The pellet was then stored in -70^0^C until use. Multidimensional Protein Identification Technology (MudPIT) was used and the data analyzed as previously described (22).

### Immunoprecipitation – Western Blotting

Approximately 10^6^ spores of the TAP tagged strains were inoculated in petri dishes containing 20 ml of Sabouraud dextrose broth at 37°C for 24 h. Mycelium that formed as a biofilm on the top was collected (∼500 mg) and was ground into fine powder in liquid nitrogen using a mortar and pestle. The ground mycelium was resuspended in 1.5 ml B250 buffer in ice cold 15ml tubes, and incubated for 10 min at 4°C, with intermittent vortexing of 30 seconds at setting 6 (Vortex Genie 2 Scientific Industries) every 2 min. The tubes containing resuspended mycelia was centrifuged at 5000 x g for 10 min at 4^0^C. 50 μl of supernatant was kept aside as input control while 750 μl was used for immunoprecipitation with HA monoclonal antibody 2-2.2.14 (Invitrogen) at 1:100 dilution and incubated on a rotating platform for 16 hours at 4^0^C. The cell lysate-Ab was then added to 50 μl of protein G Dynabeads (Invitrogen) for 6 hrs on a rotator at 4^0^C. The cells were washed with 2x with W150 buffer, and 1x with W150 buffer. The protein was eluted in 50 μl of 2x Laemmli sample buffer (Bio-rad) after heating the dynabeads at 95°C for 10 min. The input sample was also resuspended in 50 μl of 2x Laemmli sample buffer and incubated at 95^0^C for 10 min. 20 μl of input & immunoprecipitated sample was used for western blotting, which was done as described in (23). The AtrR polyclonal antibody used here has been described in (13), and was used at a 1:500 dilution, while the TAP antibody (Genscript, Inc) and the HA monoclonal antibody 2-2.2.14 (Invitrogen) were used at a 1:1000 & 1:2500 dilution, respectively. AbcG1 polyclonal antibody (24) was used at a dilution of 1:500.

### Radial growth/drug disc diffusion assay

Fresh spores of *A. fumigatus* were suspended in 1x phosphate-buffered saline (PBS) supplemented with 0.01% Tween 20 (1x PBST). The spore suspension was counted using a hemocytometer to determine the spore concentration. Spores were then appropriately diluted in 1x PBST. For the drug diffusion assay, 1×10^6^ spores were mixed with 10 ml soft agar (0.7%) and poured over 15 ml regular agar containing (1.5%) minimal medium. A paper disk was placed on the center of the plate, and 10 μl of either 1 mg/liter voriconazole was spotted onto the sterile filter paper. For the radial growth assay, ∼100 spores (in 4 μl) were spotted on minimal medium with or without the drug. The plates were incubated at 37°C and inspected for growth every 12 h.

### Real-time PCR

Reverse transcription quantitative PCR (RT-qPCR) was performed as described in reference (23), with the following modification. Cell lysates were prepared from mycelial biofilm cultures formed upon inoculating 10^6^ spores in a petri dish containing 20 ml of Sabouraud dextrose broth and grown for 24 h at 37°C under nonshaking conditions. The threshold cycle (C_T_) value of the *tef1 (Afu1g06390)* transcript was used as a normalization control.

### Fluorescent Microscopy

To visualize the localization of NcaA-GFP2-5 and AtrR-mNeonGreen, conidia were inoculated in 200 μl of minimal media on coverslip and incubated in moisture-condition at 37°C for 16 h (25). Coverslips were washed twice with 1X PBS and treated with 3 μg/ml Hoechst dye, used as nuclear stain, for 15 mins. Images were captured at 100x using Olympus fluorescent microscope BX60 controlled by iVision software (BioVision Technologies) and equipped with Hamamatsu Orca-R2 digital camera (Hamamatsu, JP). GFP filter was used for visualization of tagged proteins with excitation wavelength of 470 nm and emission wavelength of 509 nm. Excitation and emission wavelength were 359 nm and 461 nm respectively for visualization of Hoechst staining. Adobe Photoshop 2022 was used for preparing images for publication.

## Results

### Generation of AtrR-TAP fusion strain

To facilitate purification of AtrR from *A. fumigatus* cells, we constructed a fusion gene between *atrR* and a C-terminal tandem affinity purification (TAP) module that had been codon optimized for use in *Aspergillus* (18). We also fused this TAP moiety to a version of *atrR* in which its normal promoter had been replaced with the powerful *hspA* promoter (26). These constructs are diagrammed in Figure 1A. We have previously found use of the *hspA* promoter to control *atrR* expression led to overproduction of AtrR (12, 13). To test the function of these forms of *atrR*, we compared the voriconazole resistance phenotype of isogenic wt and *hspA-atrR* fusion genes with their TAP-tagged counterparts using a disk diffusion assay (Figure 1B).

**Figure 1.**
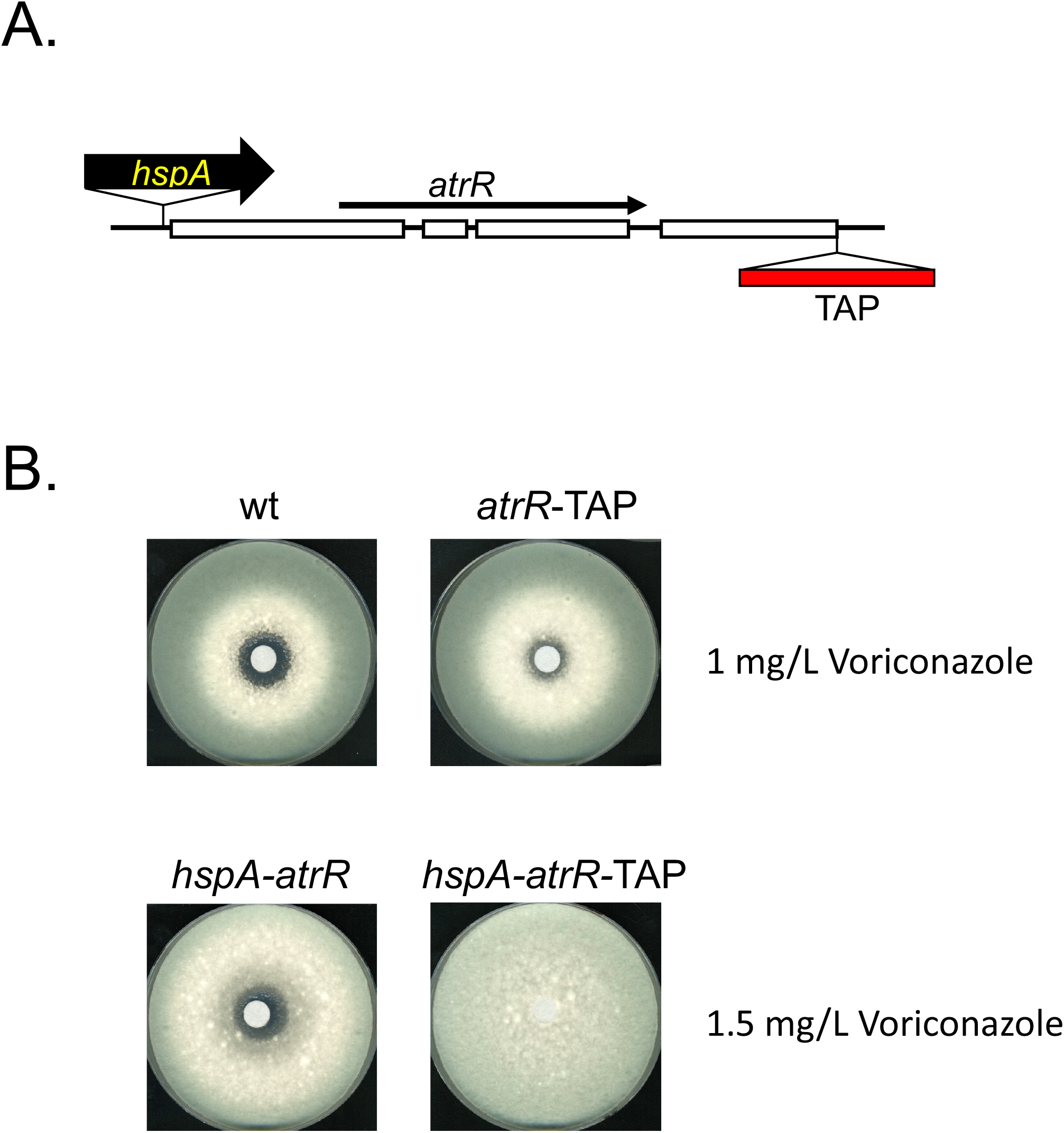
Characterization of *atrR*-TAP-expressing strains. A. The alterations made to the *atrR* locus are diagrammed. The TAP tag cassette was inserted in place of the native *atrR* stop codon and expressed using either the native *atrR* promoter or with the *A. fumigatus hspA* promoter inserted immediately upstream of the native ATG codon. Exons and introns are indicated as open bars or solid lines respectively. Transcriptional direction of *atrR* is indicated by the line. B. Voriconazole susceptibility comparison of tagged and untagged atrR alleles. An equal number of spores of each indicated strain was plated on minimal medium and allowed to dry. A sterile filter disk containing the indicated dose of voriconazole was then placed in the center of the plate. Plates were incubated at 37° C and photographed.

Introduction of the TAP tag module led to a decrease in voriconazole susceptibility in both the *atrR* and *hspA-atrR* formats. As we previously observed (12, 13) when a 3X HA epitope tag was placed at the C-terminus of AtrR, these TAP fusion proteins appeared to enhance the activity of the resulting AtrR fusion protein when compared to the wild-type factor.

### Expression of TAP-tagged forms of AtrR

Having confirmed that the introduction of the TAP moiety to the AtrR C-terminus did not prevent function of the resulting factor, we assessed expression of these protein forms compared to the wild-type protein. Transformants expressing AtrR-TAP under control of the wild-type *atrR* or *hspA* promoter were grown overnight and whole cell protein extracts prepared. We analyzed isogenic versions of these strains lacking the TAP tag as controls. Equal amounts of extracts were analyzed by western blotting using either anti-AtrR or anti-TAP antibodies.

Expression of the AtrR-TAP fusion protein led to production of the expected higher molecular mass protein when driven by the wild-type atrR promoter and the wild-type AtrR species was no longer visible (Figure 2A). Based on the relative signals of AtrR and AtrR-TAP, the TAP fusion protein appeared to be expressed at a higher level, consistent with the decreased voriconazole susceptibility in this strain. Insertion of the *spA* promoter upstream of either the wild-type *atrR* gene or the atrR-TAP allele produced higher levels of each respective AtrR form. The presence of the *hspA-atrR*-TAP fusion produced both the higher molecular mass AtrR-TAP form as well as polypeptide that was close in size to that of the untagged AtrR. We suspect this may result from proteolytic removal of the C-terminal TAP tag during this analysis. These same protein extracts were probed with the anti-TAP antibody (Figure 2B).

**Figure 2.**
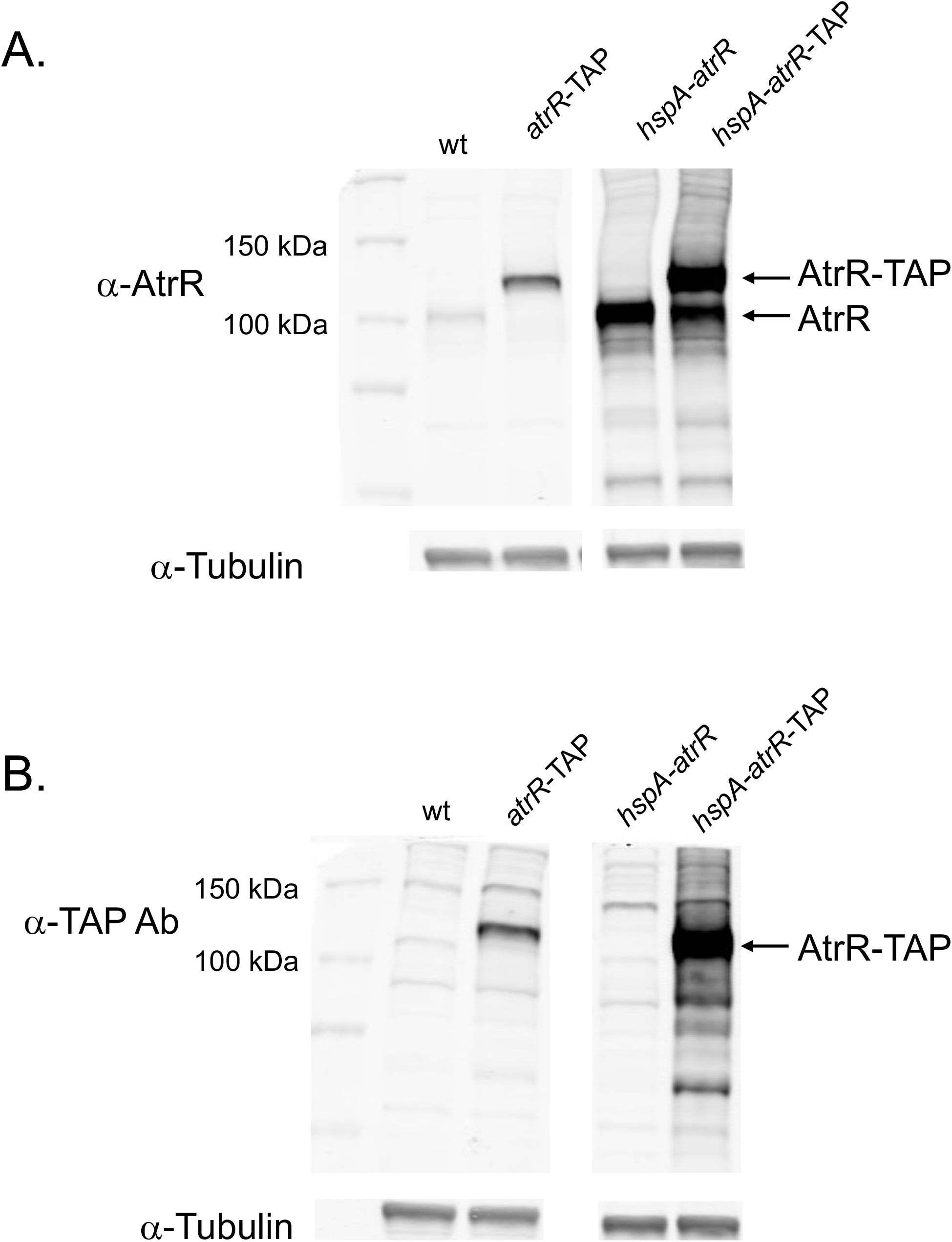
Western blot analysis of TAP-tagged *atrR* alleles. Whole cell protein extracts were prepared from *A. fumigatus* strains containing the indicated forms of the *atrR* gene. Equal amounts of protein were resolved on SDS-PAGE, transferred to nitrocellulose filters and probed with rabbit polyclonal antibodies directed against AtrR (α-AtrR) or the TAP moiety (α-TAP) or a mouse monoclonal antibody recognizing the tubulin protein (α-Tubulin). A. Use of anti-AtrR antiserum. Note the appearance of full-length AtrR in hspA-atrR-TAP strains. This is thought to be a result of proteolytic removal of the TAP tag. B. Use of the anti-TAP antibody.

A prominent polypeptide of 120 kD was seen in both strains containing the *atrR*-TAP fusion gene with the *hspA-atrR*-TAP strain producing higher levels of this protein. Some smaller polypeptides were also detected in this AtrR-TAP overproducing strain, likely as a result of proteolysis. Only background signals were detected in the absence of the inserted TAP tag.

These data suggest that AtrR-TAP accumulates (at least under these conditions) as primarily a full-length protein and that the increased levels caused by TAP fusion might contribute to the increased level of voriconazole resistance seen in strains containing the *atrR*-TAP fusion gene. To directly evaluate expression of AtrR target genes, we used western blotting to examine levels of Cyp51A and AbcG1.

We have previously described production of rabbit antibodies that can detect expression of the voriconazole target enzyme Cyp51A and the ATP-binding cassette (ABC) transporter protein AbcG1 (23). Isogenic strains containing either *atrR* or *hspA-atrR* genes with or without a TAP fusion attached were grown overnight and whole cell protein extracts prepared. These extracts were analyzed using either the anti-Cyp51A (Figure 3A) or anti-AbcG1 (Figure 3B) antibodies.

**Figure 3.**
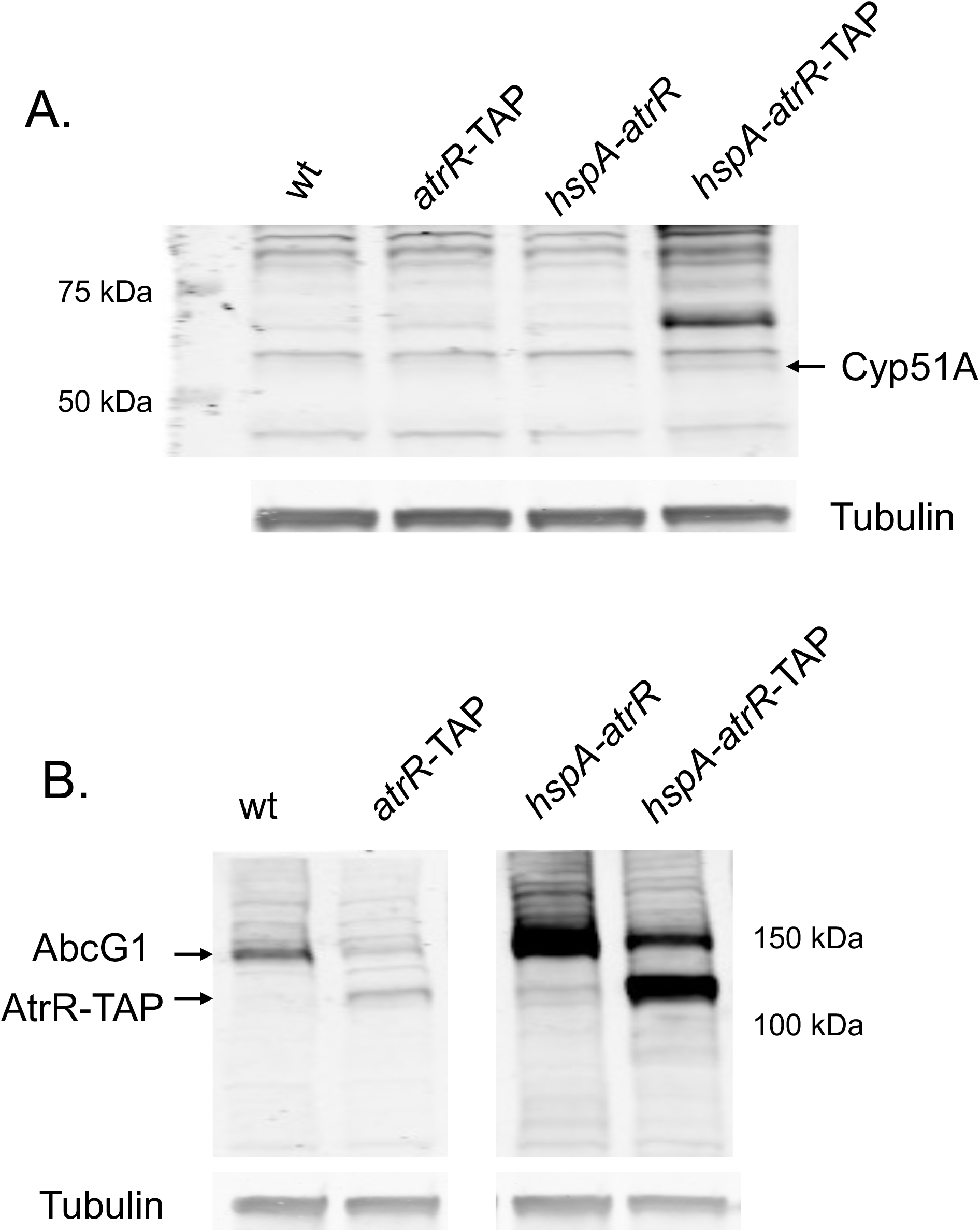
Function of TAP-tagged forms of AtrR. A. Western blot analysis of Cyp51A expression in response to the presence of the indicated forms of the *atrR* gene. Whole cell protein extracts were made from strains grown for 18 hours. Equal aliquots of each extract were electrophoresed on SDS-PAGE and then analyzed by western blotting using an anti-Cyp51A antibody (23). Location of the Cyp51A protein is indicated at the right. Tubulin was blotted as above to provide a loading control. B. Western analysis of AbcG1 expression in strains with tagged or untagged forms of AtrR. The extracts described above were analyzed by western blotting using a rabbit polyclonal antiserum directed against AbcG1. Note that the ZZ domain present in the TAP tag is able to bind the Fc chain of immunoglobulins (https://pubmed.ncbi.nlm.nih.gov/14562106/). This leads to cross-reaction with the antibodies used to detect AbcG1.

Expression of Cyp51A was only detected in the *hspA-atrR*-TAP strain. We have previously found that Cyp51A was undetectable using this western blot assay in wild-type cells but could be induced by voriconazole induction or hyperactive promoter variants such as TR34 or TR46 (8, 9). Here we were able to detect Cyp51A expression in the absence of any drug challenge in the presence of the *hspA-atrR*-TAP allele. This high basal level of Cyp51A in this strain may explain its elevated voriconazole resistance (Figure 1B).

Western blotting for AbcG1 expression in these same backgrounds produced a distinctly different result. Expression of AbcG1 was higher in the presence of the untagged alleles of *atrR* when compared to the TAP-tagged versions (Figure 3B). The presence of the *hspA* promoter led to increased AbcG1 expression when compared to the same *atrR* protein form driven by the wild-type *atrR* promoter. Note the presence of cross-reaction of the TAP fusion proteins with the rabbit primary antibody, a well-known complication of this epitope tag (27).

The differential response of *abcG1* and *cyp51A* expression to these different forms of AtrR suggests a promoter-specific effect of this transcriptional regulator. Further studies described below support this suggestion.

### Purification of AtrR-TAP

Having established that the AtrR-TAP fusion protein was able to function in vivo (albeit with some differences compared to the wild-type factor), we prepared native extracts and used standard TAP chromatographic approaches to purify this protein along with co-purifying polypeptides. We employed multidimensional protein identification technology (MudPIT) to detect these co-purifying proteins and found many different candidates (16). Here we will focus on a single protein that co-purified with AtrR-TAP and was 2X more abundant in AtrR-TAP fractions from the *hspA*-driven fusion gene than when produced from the wild-type *atrR* promoter. This protein was designated <u>n</u>uclear <u>c</u>o<u>a</u>ctivator of <u>A</u>trR or NcaA. NcaA is encoded by the gene *AFUA_6g08010*. While the predicted polypeptide produced by *ncaA* represented a previously undescribed protein, although present in most fungal species, we provide evidence supporting the NcaA designation below.

### NcaA interacts with AtrR in vivo

To further support our identification of NcaA as an interactor with AtrR, we carried out a coimmunoprecipitation analysis using epitope-tagged forms of these two proteins. The *hspA-atrR*-TAP-containing strain was used as to allow facile identification of AtrR and a 3X HA-tagged form of ncaA was introduced into this strain. Isogenic *hspA-atrR*-TAP strains either containing or lacking the *ncaA*-3X HA allele were grown overnight, native protein extracts prepared and the NcaA-3X HA protein recovered by immunoprecipitation with anti-mouse HA antibody. These anti-HA immunoprecipitates were electrophoresed on SDS-PAGE and then analyzed by western blotting using either anti-AtrR or anti-HA antibodies (Figure 4).

**Figure 4.**
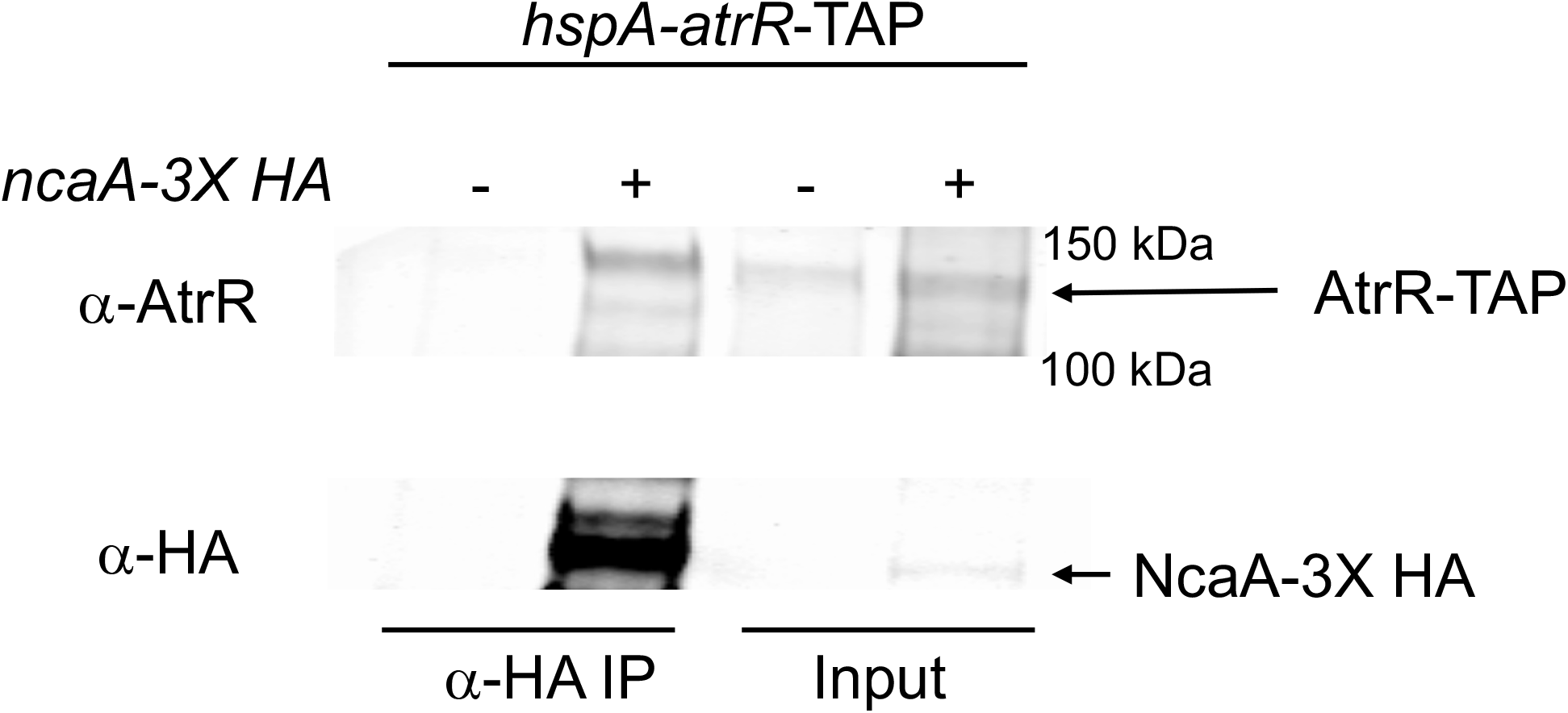
Association of NcaA and AtrR in vivo. A strain expressing AtrR-TAP and either containing (+) or lacking (-) an *ncaA*-3X HA fusion gene was grown for 18 hours in YG medium. Whole cell protein extracts were prepared under native lysis conditions and used for immunoprecipitation with an anti-HA antibody. Samples of the native total lysate were retained to confirm the presence of each protein (Input). HA-immunoprecipitates were recovered and run in parallel followed by western blotting with either anti-AtrR or anti-HA antibodies. The location of each protein is indicated at the right hand side.

Only when both the NcaA-3X HA-tagged allele and the AtrR-TAP fusion were present was coimmunoprecipitation seen. Expression of only the AtrR-TAP fusion protein did not show any evidence for nonspecific recovery of this factor by the anti-HA antibody. These data support the view that AtrR and NcaA associate in vivo.

### NcaA is required for normal voriconazole resistance

We generated a strain lacking the *ncaA* coding sequence using a CRISPR-based gene deletion strategy (19). Spores were produced from isogenic wild-type and *ncaAΔ* strains and plated on minimal media. A filter disk containing different concentrations of voriconazole was placed in the center of these spores and the resulting plate incubated to allow growth of the cells. The distance from the disk at which growth stopped (zone of inhibition) was measured. These experiments were performed on three independent isolates of *ncaAΔ* (Figure 5).

**Figure 5.**
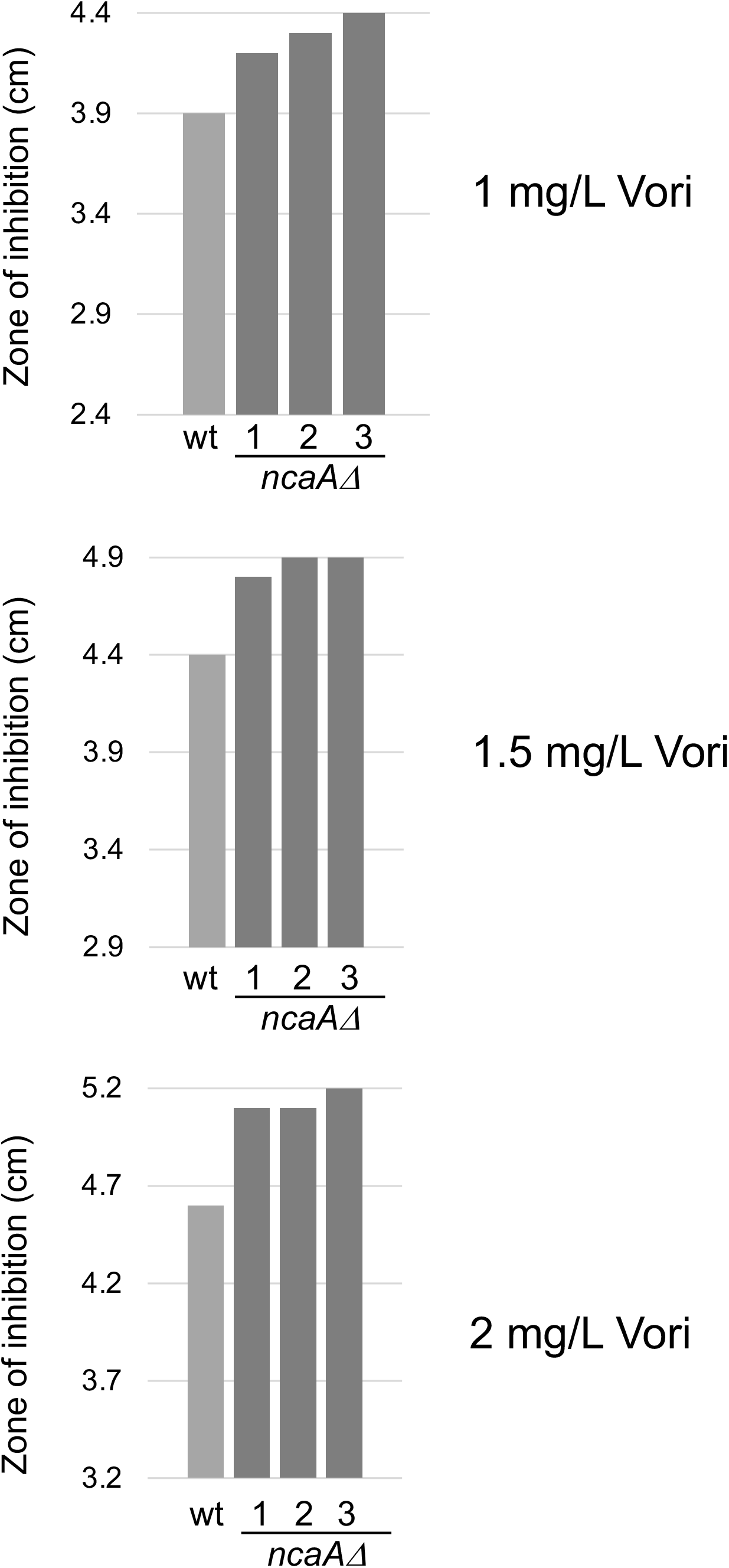
Loss of *ncaA* increased voriconazole susceptibility. Wild-type AfS35 cells and isogenic ncaAD mutants were grown and analyzed for voriconazole susceptibility using a disk diffusion assay as shown above. Different doses of voriconazole were applied to each disk and the plates incubated at 37°C. The distance from the edge of the disk to the beginning of growth (zone of inhibition) was measured for 3 independent isolates of the *ncaAΔ* strain.

Loss of *ncaA* caused a modest but highly reproducible increase in voriconazole susceptibility. These data are consistent with NcaA playing a positive role in conferring voriconazole tolerance.

### NcaA is required for azole-induced expression of an AtrR target gene

To probe the requirement for NcaA in AtrR-dependent gene regulation, isogenic wild-type and *ncaAΔ* strains were grown overnight in the presence or absence of sublethal doses of voriconazole. These cultures were harvested, total RNA prepared and analyzed by quantitative reverse transcription followed by PCR analysis of three different mRNA species corresponding to known AtrR target genes. We used the *abcG1, cyp51A* and *atrR* genes as representative AtrR-controlled genes.

Transcription of *abcG1* was induced by approximately 3.5-fold in the presence of voriconazole in wild-type cells but this was reduced to approximately 2-fold in *ncaAΔ* strains. Loss of *ncaA* failed to impact expression of either *cyp51A* or *atrR*. This modest reduction in *abcG1* induction in the presence of voriconazole is consistent with the level of increased susceptibility seen earlier (Figure 5).

### NcaA is localized to the nucleus

Based on its copurification with AtrR, we suspected that NcaA would be localized to the nucleus. We believed AtrR would be a nuclear factor based on its clear role as a regulator of gene expression and the ability to detect AtrR bound to its DNA target sites. To test these predictions, we constructed C-terminal fusion genes between *ncaA* and green fluorescent protein (GFP) as well as *atrR* with A. fumigatus codon-optimized mNeonGreen (mNG). Strains containing either the *ncaA*-GFP or *atrR*-mNG fusion genes were grown overnight and then visualized by microscopy (Figure 6).

**Figure 6.**
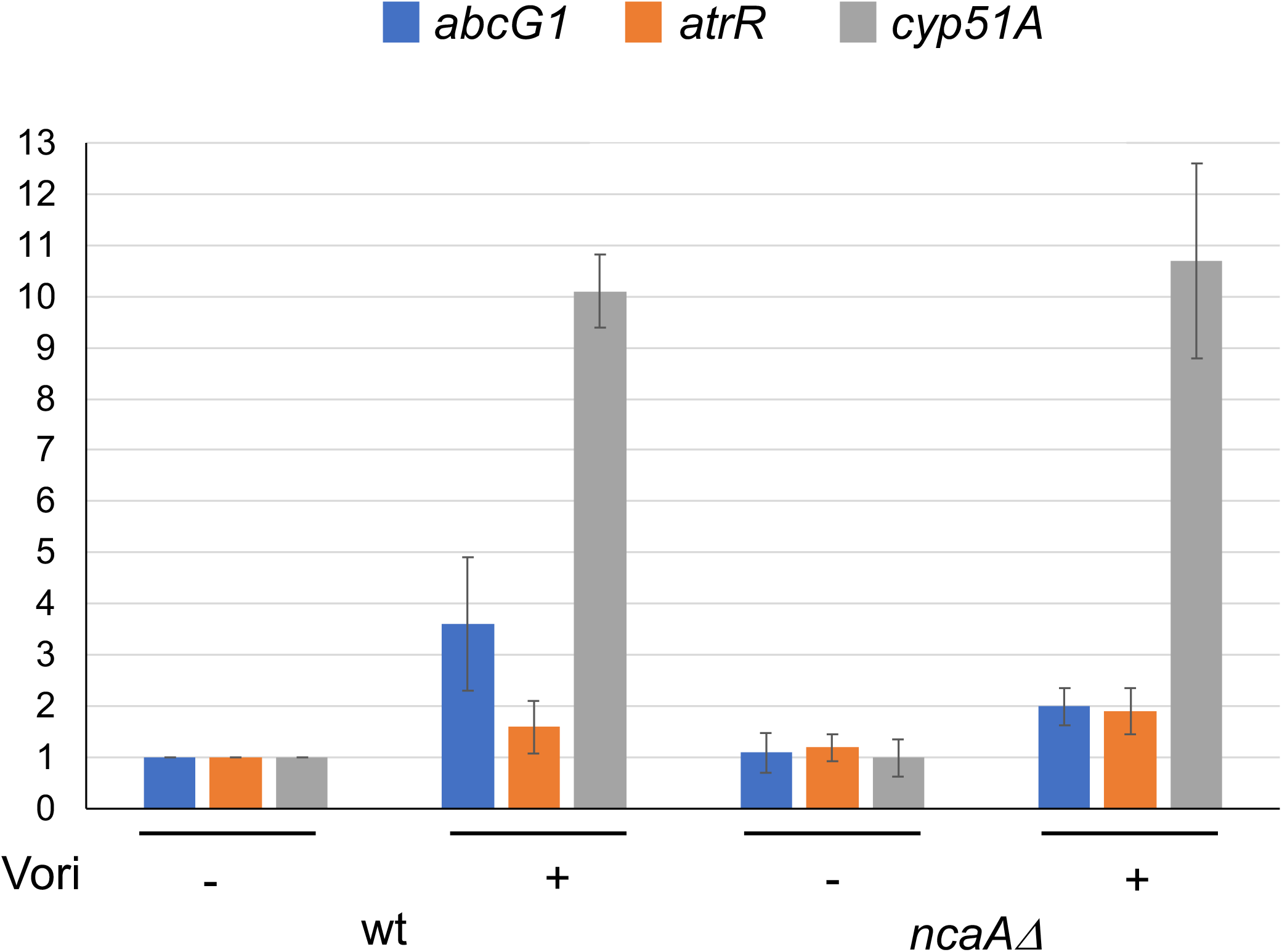
Absence of *ncaA* prevents normal voriconazole induction of *abcG1* expression. Isogenic wild-type and *ncaAΔ* strains were grown in presence (+) or absence (-) of voriconazole. RNA was prepared from each culture and steady-state mRNA levels of the indicated genes measured using qRT-PCR.

**Figure 7.**
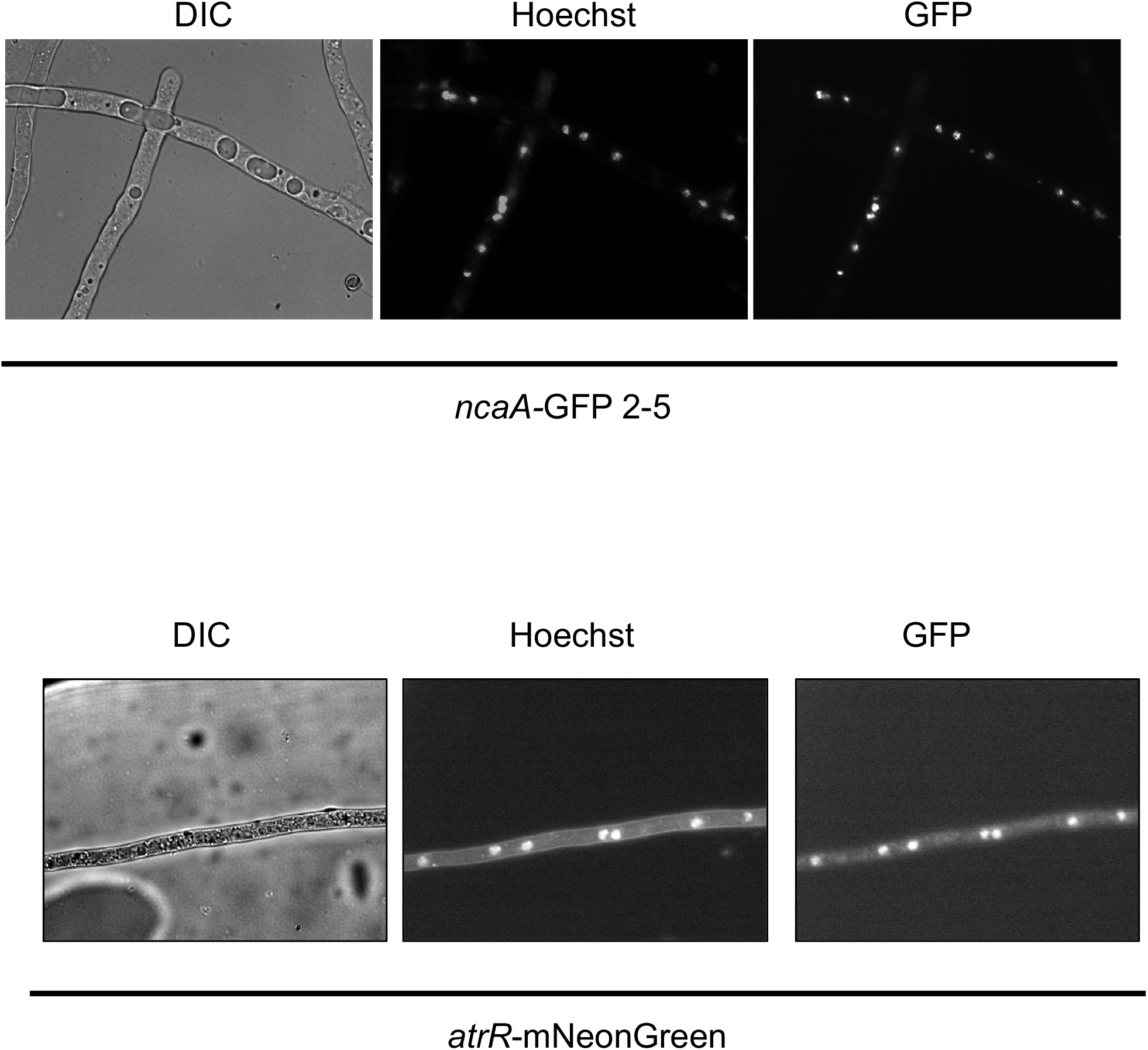
NcaA and AtrR are both localized to the fungal nucleus. A. fumigatus strains expressing either a ncaA-GFP fusion protein (top panel) or an atrR-mNeonGreen fluorescent protein (bottom panel) were grown overnight and then analyzed by light microscopy. Cells were visualized by Nomarski optics (DIC), nuclei detected by staining with Hoechst dye. Nuclei and GFP were visualized with appropriate filters.

The NcaA-GFP fusion protein was found to be localized to the nucleus in *A. fumigatus* hyphae. Similarly, AtrR-mNG was also found in the nucleus. We confirmed the identity of the nuclear compartment by staining nuclear DNA with Hoechst dye. NcaA association with AtrR is likely to involve their association within this organelle.

## Discussion

AtrR is a major determinant of azole resistance in *A. fumigatus* but little is known of how this factor is regulated (12, 13). To identify proteins that may act to modulate AtrR function, we prepared and purified a TAP-tagged version of this transcription factor. We were able to detect a number of different proteins that copurified with AtrR-TAP and have focused here on a previously uncharacterized protein we have designated NcaA. NcaA is a novel protein with no obvious conserved structural domains. Analysis of the sequence of NcaA predicts a central region with coiled coil domains (unpublished data) flanked by more disordered and flexible regions. Most *Aspergillus* species contain an orthologue of NcaA suggesting that the function of this protein must be conserved across these related organisms.

Our data support a role for NcaA in transcriptional activation based on two different assays. First, loss of NcaA produced an increase in voriconazole susceptibility across a range of drug concentrations (Figure 4). Second, a gene-specific defect in drug induction was seen for the AtrR target gene *abcG1*. Two other AtrR target genes were unaffected by loss of NcaA. The observation that drug induction of both *atrR* itself and the azole drug target-encoding gene *cyp51A* was unaffected is a likely a central factor in determining the resulting voriconazole susceptibility of the *ncaAΔ* strain. Our previous analyses of both the *abcG1* and *cyp51A* promoters may help explain the differential effects of the *ncaAΔ* allele on these two genes. Transcription of *abcG1* is highly dependent on AtrR activity while expression of *cyp51A* involves both AtrR as well as the key sterol regulatory transcription factor SrbA (10, 11, 14, 28, 29). NcaA may not be involved in AtrR-dependent activation at the *cyp51A* promoter or the partially compromised phenotype triggered by the *ncaAΔ* allele may be suppressed by the presence of normal SrbA.

This gene specific activation by AtrR is also seen in the characterization of the *atrR*-TAP allele. Cyp51A could be detected by western blotting only in cells containing the *hspA-atrR*-TAP allele, not in cells expressing *atrR*-TAP from the native *atrR* promoter or *hspA-atrR*-containing strains. In contrast to the response of the *cyp51A* promoter, expression of the *atrR*-TAP fusion protein was generally less effective at driving transcription of *abcG1*. These data argue that while the presence of the TAP tag at the C-terminus of AtrR prevents normal gene activation at the *abcG1* promoter, this same recombinant protein appears to be a more effective activator of *cyp51A*. We have previously documented that a 3X HA tag at the C-terminus of AtrR behaves as a hypermorphic (activated) allele of *atrR* (13). Interestingly, the AtrR-3X HA seemed to be a better inducer of *abcG1* expression than *cyp51A*. The variable effects of these different AtrR fusion proteins suggests that the contacts made by this factor at different promoter are unique.

The nonidentical responses of AtrR target promoters to different AtrR fusion proteins illustrates the complexity of transcriptional activation by this factor. This is also likely to contribute to the phenotype caused by loss of NcaA. Purification of AtrR-TAP yields a population of complexes that represent an average of proteins associated with AtrR. Promoter-specific complexes may be formed that are recovered together during our purification. Further analyses are required to determine how the various factors that associate with AtrR contribute to the function of this protein and if these contribute equally at the various target promoters responsive to AtrR.

## Acknowledgements

We thank Drs. Damian Krysan, Paul Bowyer and Michael Bromley for helpful discussions and Bastian Joehnk and Gerhard Braus for providing the codon-optimized TAP clone. This work was supported by NIH grant AI143198.

